# Voluntary wheel running improves outcomes in an early life stress-induced model of urologic chronic pelvic pain syndrome in male mice

**DOI:** 10.1101/2020.07.20.212480

**Authors:** Isabella M. Fuentes, Brittni M. Jones, Aaron D. Brake, Angela N. Pierce, Olivia C. Eller, Rachel M. Supple, Douglas E. Wright, Julie A. Christianson

## Abstract

Patients with a history of early life stress (ELS) exposure have an increased risk of developing chronic pain and mood disorders later in life. The severity of ELS in patients with urologic chronic pelvic pain syndrome (UCPPS) is directly correlated with symptom severity and increased comorbidity, and is inversely related to likelihood of improvement. Voluntary exercise improves chronic pain symptoms and our group and others have shown that voluntary wheel running can improve outcomes in stress-induced UCPPS models, suggesting that exercise may negate some of the outcomes associated with ELS. Here we provide further evidence that voluntary wheel running can attenuate increased perigenital mechanical sensitivity, bladder output, and mast cell degranulation in the bladder and prostate in male mice that underwent neonatal maternal separation (NMS). Sedentary male NMS mice had reduced serum corticosterone, which was not impacted by voluntary wheel running, although stress-related regulatory gene expression in the hypothalamus and hippocampus was significantly increased following exercise. Neurogenesis in the dentate gyrus of the hippocampus was diminished in sedentary NMS mice and significantly increased in both exercised naïve and NMS mice. Sucrose consumption increased in exercised naïve but not NMS mice, and anxiety behaviors measured on an elevated plus maze were increased following exercise. Together these data suggest that voluntary wheel running is sufficient to normalize many of the UCPPS-related outcomes resulting from NMS. Exercise also increased hippocampal neurogenesis and stress-related gene expression within the hypothalamic-pituitary-adrenal axis, further supporting exercise as a non-pharmacological intervention for attenuating outcomes related to ELS exposure.

## Introduction

Chronic prostatitis/chronic pelvic pain syndrome (CP/CPPS) is the most common urological diagnosis for men under the age of 50 and affects approximately 8-11.5% of men in the United States (1, 2). The main symptoms of CP/CPPS include pain and discomfort within the urogenital organs, with or without increased urgency and voiding, that often worsen on ejaculation (1). The symptoms of CP/CPPS are often indistinguishable with those of interstitial cystitis/painful bladder syndrome (IC/PBS). Early life stress (ELS) exposure is reported at a higher rate among patients with urologic chronic pelvic pain syndrome (UCPPS), which encompasses both CP/CPPS and IC/PBS (1). In a multi-site study of UCPPS patients, the severity of ELS was correlated with greater widespread pain and negative mood, and was inversely associated with likelihood of symptomatic improvement (3). Neuroimaging studies in the same cohort of UCPPS patients revealed significant changes in gray matter volume, neurochemical concentration, and functional connectivity that are correlated with widespreadness of pain and comorbid mood disorder (4–9). Further analysis showed both sex- and ELS-dependent effects in discreet pain-processing regions (10), suggesting that ELS may specifically predispose patients to greater symptom burden and comorbidity.

Exposure to ELS significantly impacts the development and regulation of the hypothalamic-pituitary-adrenal (HPA) axis, which regulates the response to stress. The hippocampus, which provides inhibitory regulation to the hypothalamus, is negatively impacted by ELS exposure, as evidenced by decreased gray matter volume (11–16). Corticotropin-releasing factor (CRF) is a major signaling hormone involved in HPA axis activation and regulation. CRF is released by the hypothalamus and causes downstream production of glucocorticoids and local activation of resident immune cells in peripheral tissues. Mast cells express CRF receptor 1 (CRF_1_) and can become degranulated and/or release pro-inflammatory mediators upon activation (17). Increased mast cell tryptase levels have been reported in expressed prostatic secretions from CP/CPPS patients and biopsies have also shown histological evidence of increased mast cell degranulation (18, 19).

Voluntary exercise has been shown to attenuate molecular and behavioral changes associated with a dysfunctional HPA axis (20) and significantly improves symptom severity in patients with IBS (21–26), fibromyalgia (27–29), and depression and/or anxiety (30–32). Voluntary wheel running has a significant impact on increasing hippocampal neurogenesis and brain-derived neurotrophic factor (BDNF) expression, and preventing depression-like behaviors in rodents (33–36). We and others have shown that voluntary exercise can normalize abnormal hippocampal gene expression resulting from neonatal maternal separation (NMS) in rodents, thereby reinstating proper descending limbic control of the HPA axis (37, 38). The goal of the current study was to gain novel insight on the mechanisms underlying comorbid pelvic pain and mood disorders following early life stress and the potential therapeutic impact of voluntary wheel running for mitigating these outcomes. We report behavioral measures of perigenital sensitivity and mood disorder, as well as histological and molecular evidence of altered HPA axis regulation and hippocampal neurogenesis.

## Methods

### Animals

All experiments in this study were performed on male C57Bl/6 mice that were born from pregnant dams (Charles River, Wilmington, MA) delivered to the Research Support Facility at the University of Kansas Medical Center during the third week of gestation. Mice were housed in a climate-controlled room on a 12-h light cycle from 600 to 1800 h and received water and food *ad libitum*. Animal use protocols conformed to NIH guidelines and were approved by the University of Kansas Medical Center Institutional Animal Care and Use Committee and the Committee for Research and Ethical Issues of IASP.

### Neonatal maternal separation

Beginning on postnatal day 1 (P1, date of birth was considered P0), pups were removed daily from their home cages for 180 minutes (1100 to 1400 hours) and placed as a litter, with a small amount of home bedding material, into a clean glass beaker and held at 34°C and 50% humidity as previously described (39). NMS mice underwent daily separation from P1 through P21 and were weaned at P22. Naïve mice were born in-house and remained undisturbed in their home cages, with the exception of routine animal husbandry, until weaning at P22. Four separate cohorts of NMS mice were used in this study and each cohort was compared to a corresponding naïve group of mice that were born, housed, and weaned during the same time frame to avoid potential complications arising from variations in prenatal shipping conditions, housing environment, and investigator handling. Although only male mice are reported on here, all litters were maintained with both male and female pups until weaning.

### Voluntary wheel running

At four weeks of age, male naïve and NMS mice were either pair-housed in cages equipped with a stainless-steel running wheel (Ex mice; STARR Life Sciences Corp, Oakmont, PA) or remained group housed (2-4/cage) in standard cages with IACUC-compliant enrichment standards and no running wheel (Sed mice). Running distance was measured by STARR Life Sciences VitalView Acquisition and Analysis software version 1.1.

### Perigenital mechanical withdrawal threshold measurement

Mice were acclimated to the testing room for two days prior to the assessment of perigenital mechanical threshold testing. On the day of testing, naïve and NMS mice were placed beneath individual clear plastic chambers (11×5×3.5cm) on a wire mesh screen elevated 55 cm above a table and allowed to acclimate for 1 hour prior to testing. The experimenter was blinded to the group status of the mice. The up-down method was performed to test mechanical sensitivity using a standard set of Semmes-Weinstein monofilaments (1.65, 2.36, 2.83, 3.22, 3.61, 4.08, 4.31, 4.74g; Stoelting, Wood Dale, IL) (40). Beginning with the 3.22g monofilament, mice received a single application to the scrotum, avoiding the midline. A negative response was followed by the next larger filament and a positive response (considered a jerk or licking) was followed by the next smaller filament. The experimenter continued to move up or down the series, depending on the previously elicited response, for an additional four applications after the first positive response was observed for a minimum of five or a maximum of nine total monofilament applications. The value in log units of the final monofilament applied in the trial series was used to calculate a 50% g threshold for each mouse and group means were determined as previously described (41).

### Micturition analysis

Surfaces were cleaned with 70% ethanol and wiped dry. Naïve and NMS mice were acclimated to the testing room in their home cages for 30 minutes prior to micturition pattern data collection. Mice were singly confined to a sheet of filter paper (11 × 5 × 3.5 cm) for 1 hour using an inverted Micro-Isolator cage bottom (Lab Products, Seaford, DE). At the end of the testing period, the filter paper was left to dry while mice were returned to home cages and the animal facility. Once the filter papers were dry, fecal pellets were counted and urine spots were visualized using UV light. The total area (cm^2^) and number of urine spots were recorded.

### Sucrose preference testing

Mice were individually housed in BioDAQ Liquid Choice Unplugged Allentown cages (Biological Data Acquisition, New Brunswick, NJ) equipped with two Polysulfone BioDAQ drinking bottles. Following a 24-hour acclimatization period during which both bottles contained standard drinking water, one bottle was filled with 1% sucrose diluted in drinking water and fluid volume levels were recorded and the position of the bottles interchanged daily for the duration of the experiment. The total volume of 1% sucrose consumed, as well as the percentage of 1% sucrose consumed was calculated on a daily basis.

### Elevated plus maze testing

Anxiety-like behaviors were measured in mice as previously described by Walf and Frye (42). Mice were placed in the center of an elevated (60 cm above the floor) plus-maze consisting of two open and two closed arms opposite each other in a plus-shaped formation. All mice were tested within the first 7 hours of the light cycle and placed into the center of the maze facing the same open arm. Mice were allowed to freely explore the maze for a testing period of 5 minutes while a digital camera captured the activity from above. The number of entries into and the time spent in open and closed arms were recorded by an observer at the time of testing in the event a video recording was unusable. Blinded scorers reviewed the recorded test sessions and were able to record the number of entries and time spent in each arm more accurately.

### Serum corticosterone ELISA

Trunk blood was collected at the time of sacrifice during the early half of the light-cycle (0800-1100 h), allowed to clot for 1 hour on ice, and centrifuged at 12,000 rpm for 10 minutes held at 2°C. Serum (clear supernatant) was collected and stored at −80°C until analysis. Serum corticosterone (CORT) was quantified using ELISA kit according to manufacturer’s instructions (ALPCO, Salem, NH).

### Acidified toluidine blue mast cell staining

Mice were overdosed with inhaled isoflurane (>5%) and intracardially perfused with ice cold 4% paraformaldehyde. Urinary bladders and prostates were removed, post-fixed in paraformaldehyde at room temperature for 1 hour, cryoprotected in 30% sucrose in 1x PBS at 4°C overnight, and frozen in a heptane bath on dry ice. Tissue was transversely cut into 10 μm-thick (bladder) and 7 μm-thick (prostate) cryosections using a cryostat held at −22°C. Cryosections were then stained with acidified toluidine blue to visualize mast cells as previously described (39). The percentage of degranulated/total mast cells was calculated in 8 separate, non-serial sections spanning the length of each tissue.

### mRNA extraction and qRT-PCR

Mice were overdosed with inhaled isoflurane (>5%). Brains were removed and frozen on dry ice. Hypothalamus, amygdala, and hippocampus were dissected, immediately snap frozen in liquid nitrogen, and stored at −80°C. Total RNA was isolated from dissected tissues using an AllPrep DNA/RNA/Protein Mini kit (Qiagen, Valencia, CA). The concentration and purity were determined using a 2100 Bioanalyzer (Agilent Technologies, Santa Clara, CA) and cDNA was synthesized from total RNA using the iScript cDNA Synthesis Kit (Bio-Rad, Hercules, CA). Quantitative RT-PCR was performed using SsoAdvanced SYBR Green Supermix (Bio-Rad) and a Bio-Rad CFX Connect Real-Time PCR Detection System with indicated 20μM primers (Integrated DNA Technologies, Coralville, IA). GAPDH was used as a housekeeping gene. Forward and reverse primer sequences are reported in Table 1.

**Table 1.**
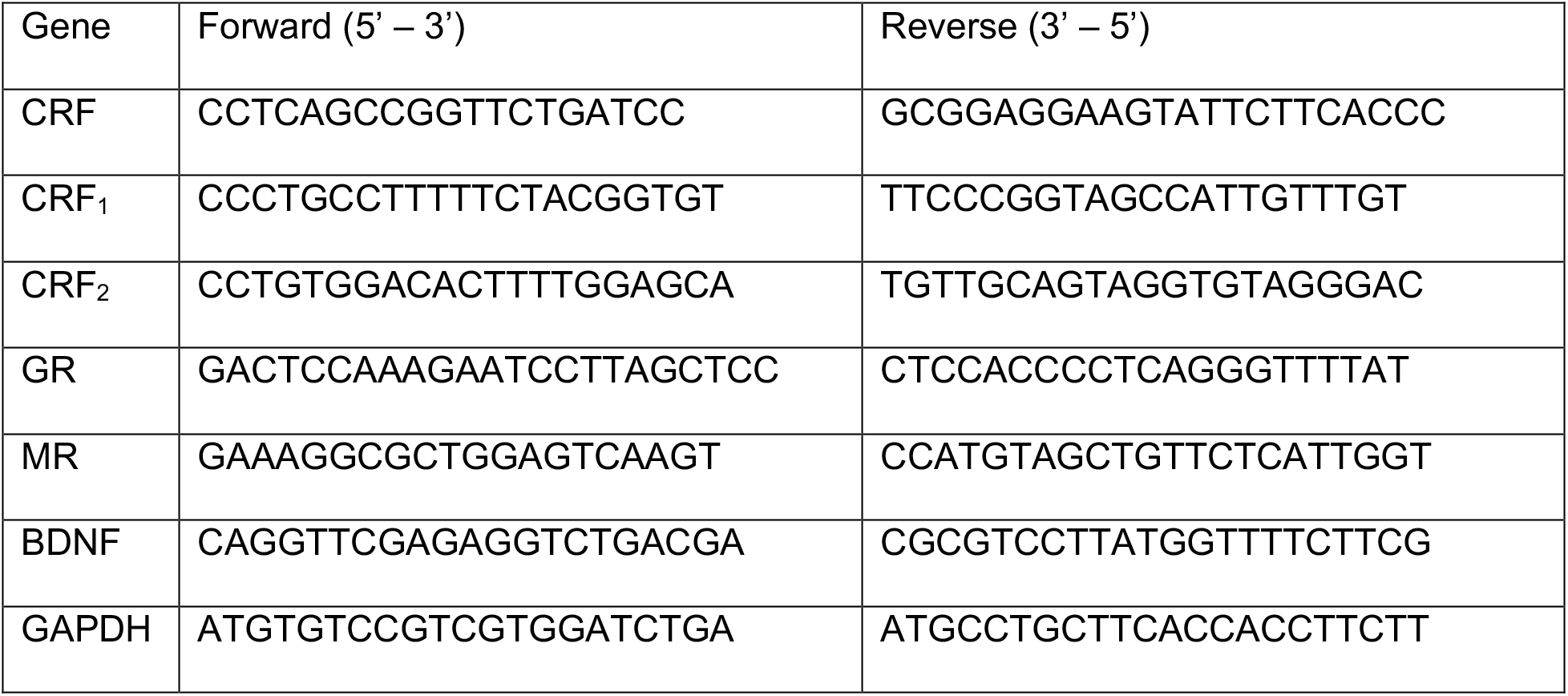
Forward and reverse primers for genes analyzed using real-time PCR in the hypothalamus, amygdala, and hippocampus.

### Intraperitoneal injection of BrdU and immunohistochemistry

Mice were given two intraperitoneal injections of 10 mg/kg of BrdU (ab 142567, Abcam, Cambridge, MA), six hours apart. Mice were sacrificed 24 hours after the first injection by inhaled isoflurane overdose (>5%) and intracardially perfused with ice cold 4% paraformaldehyde. Whole brains were removed and embedded in 10% gelatin. Trimmed blocks were post-fixed in 4% paraformaldehyde at 4°C for 2 hours on a rocker, sunk in 30% sucrose solution overnight, and stored at 4°C until cut in the coronal plane on a sliding microtome. Coronal sections, 40 μm-thick, were collected in 0.1 M phosphate buffer (PB), pH 7.4, and stored at 4°C. Once the dentate gyrus was discernable in a section, every 6^th^ section was collected in a series spanning the width of the hippocampus (dorsal to ventral).

Free floating sections underwent pretreatment as previously described (Lagace et al, 2010). Nonspecific staining was blocked by incubation with 3% normal donkey serum (NDS) in 0.1% Triton X-100 in 1X PBS for 60 minutes. Sections were then incubated in both rat anti-BrdU (1:500; Abcam) and rabbit anti-NeuN (1:1000; Abcam) in 3% NDS (vol/vol) in 0.1% Tween-20 in 1X PBS overnight at 4°C. The following day, sections were incubated with both donkey anti-rat Cy3 (1:800; Jackson Immuno Research) and Alexa Fluor-488 donkey anti-rabbit (1:500; ThermoFisher) in 1.5% NDS (vol/vol) for 60 minutes. Sections were then counterstained with Hoescht (1:3000; ThermoFisher) diluted in 0.1 M PB for 5 minutes at room temperature, then washed three times with 0.1 M PB before being mounted onto Superfrost slides (VWR, Radnor, PA) with 0.1% gelatin and coverslipped with Prolong Gold antifade reagent (Invitrogen; P36930).

### Visualization and quantification of hippocampal dentate gyrus neurons

From each mouse, 3-4 sections were examined and quantified if the dentate gyrus of the hippocampus had 2 granular cell layers present that had not yet connected. From each section, 3 images were taken: at the point of convergence of the 2 granular cell layers, the middle of the dentate with the 2 layers present, and one at the end of the dentate. The focus of these images was on the subgranular zone (SGZ) because it has been shown to give rise to new neurons that migrate to the granule cell layer where they become mature neurons (43). Therefore, NeuN labelling indicated the granule cell layer and BrdU labelling was confined to immature neurons in the subgranular zone.

The quantification of BrdU labeled neurons was performed by an examiner blinded to the treatment groups. Sections were viewed using a Leica TCS SPE II confocal microscope (Leica, Wetzlar, Germany) at 40x magnification. 30μm-thick optical sections were captured as a 4-image z-stack through the depth of the section. The number of BrdU positive cells was counted in the top and bottom image of the z-stack and then averaged. This was repeated for all 3 images captured from the same hippocampus and the average from each was added together. The total number of BrdU-positive cells was averaged for each mouse and averaged again within each experimental group.

### Statistical analysis

The timeline for the experiment is shown in Figure 1A with all outcomes being evaluated at 8 weeks of age, with the exception of weekly running distances. Calculations were made using Microsoft Excel and a Shapiro-Wilk normality test was performed to determine normality of the data. Perigenital mechanical withdrawal thresholds were log transformed and micturition data were rank transformed prior to statistical analysis. Data were analyzed using Student’s *t*-test or 2-way (with or without repeated measures) analysis of variance (ANOVA) followed by Fisher’s least significant difference or Bonferroni’s posttest (IBM SPSS Statistics, IBM Corporation, Armonk, NY; GraphPad Prism 8, GraphPad Software, La Jolla, CA), as denoted in the manuscript. All data are expressed as mean ± SEM. A *p* value of less than 0.05 was considered significant.

**Figure 1.**
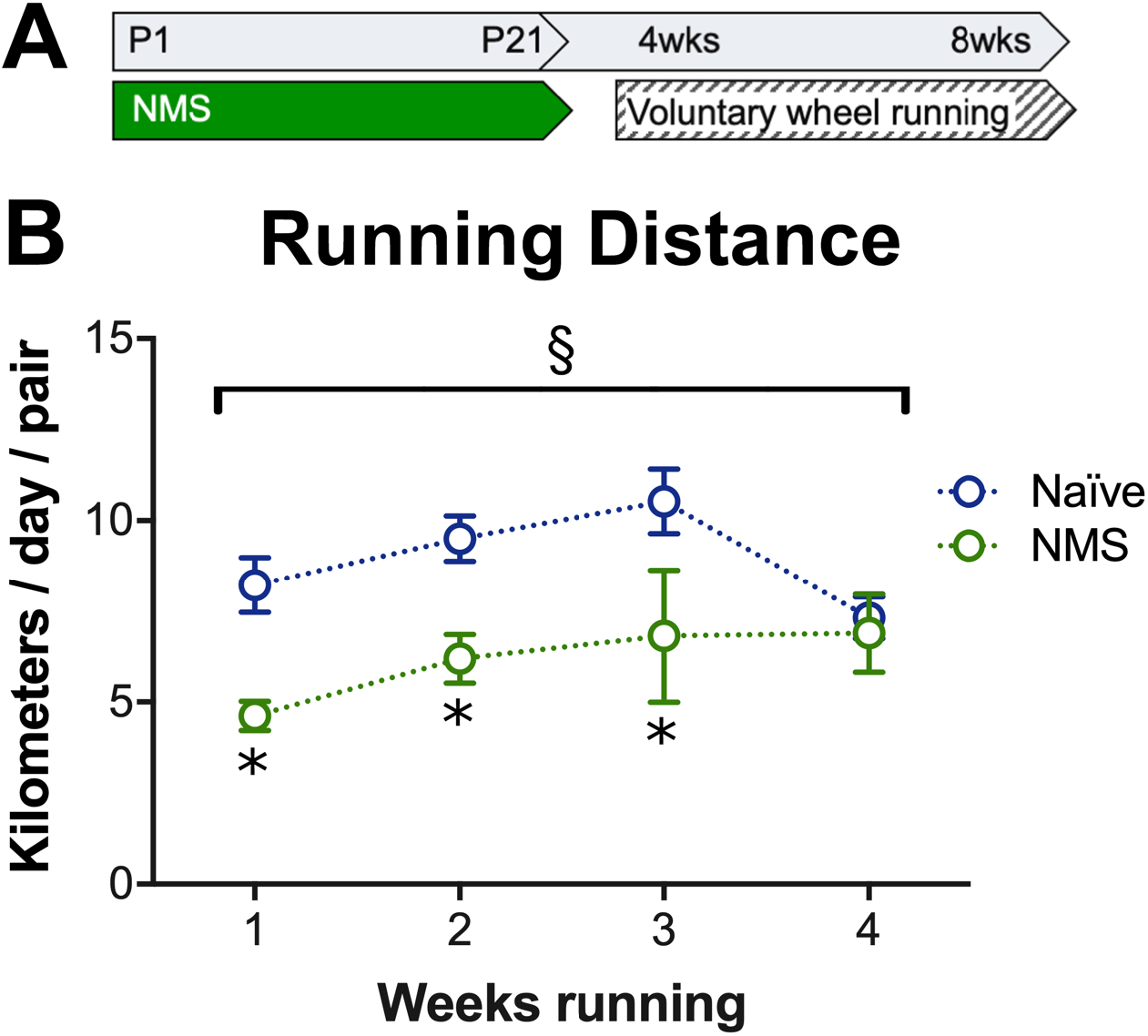
Male mice in this study underwent neonatal maternal separation (NMS) and or pair-housing in cages equipped with running wheels. **A)** A schematic represents the time line of NMS and voluntary wheel running prior to behavioral and *in vitro* assessment at 8 weeks of age. **B)** NMS had a significant impact on running wheel activity, such that NMS pairs maintained a significantly lower running distance compared to naïve mice for the first three weeks of wheel running. Bracket indicates a significant effect of NMS (§, *p*=0.0002), two-way ANOVA, \**p*<0.05, Fisher’s LSD posttest. n=6 pairs (all groups).

## Results

### Voluntary wheel running distance was shorter in male NMS mice

We previously reported that both female (38) and male (44) NMS mice ran shorter distances than naïve mice when given free access to a running wheel in their home cage. Here, we evaluated weekly cumulative running distances in pair-housed male naïve and NMS mice and, again, observed a significant decrease in running distance in NMS mice (Figure 1B). This was most prominent between 4-7 weeks of age and distances were not significantly different during the last week of observation.

### Voluntary wheel running prevented NMS-induced perigenital hypersensitivity

Consistent with our previous studies (39, 45), NMS induced significant perigenital hypersensitivity (Figure 2). Sedentary NMS mice exhibited significantly lower withdrawal thresholds in response to mechanical, monofilament stimulation to the scrotum compared to their naïve-Sed counterparts. Access to a running wheel prevented NMS-induced allodynia, such that naïve-Ex and NMS-Ex males showed similar withdrawal thresholds that did not significantly differ from naïve-Sed males.

**Figure 2.**
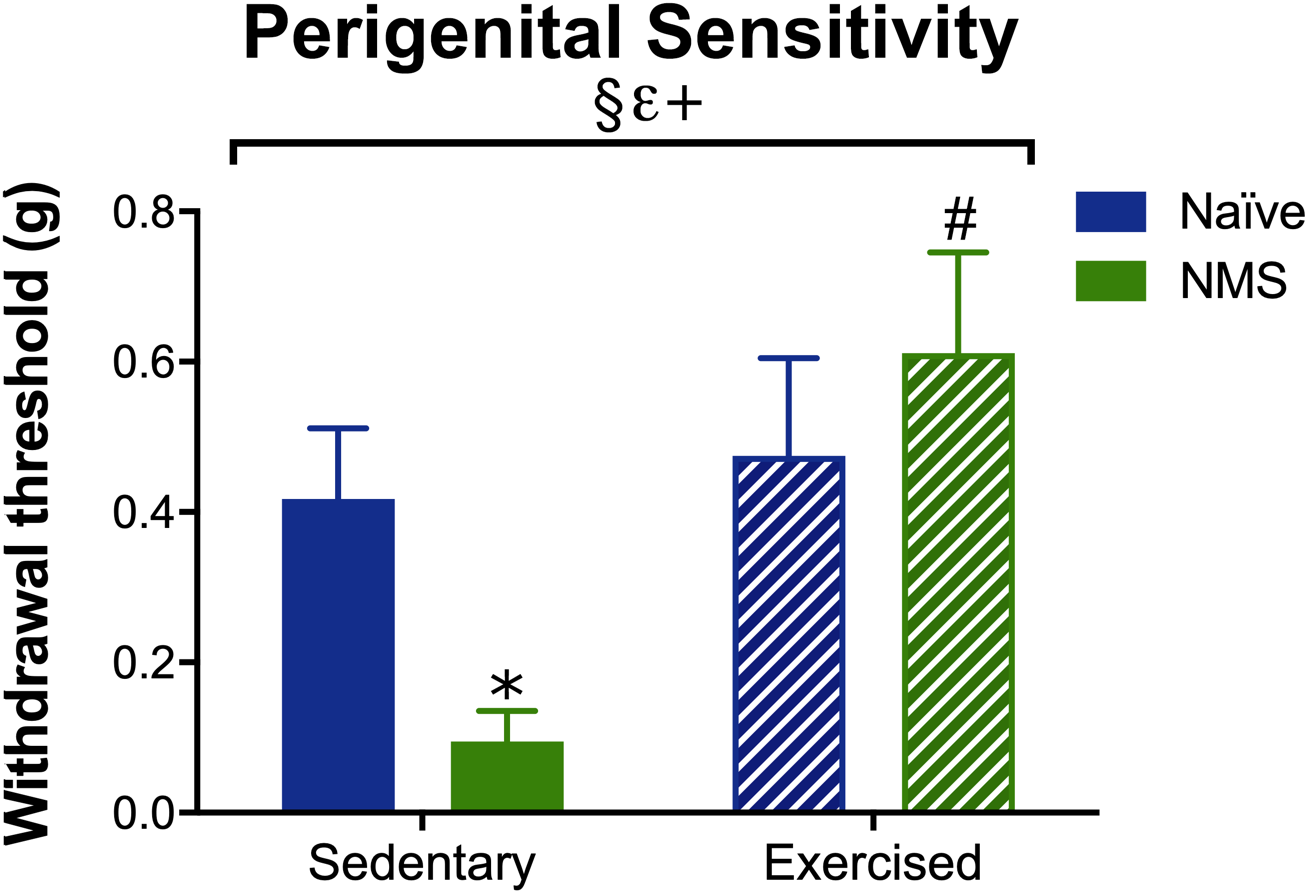
Mechanical withdrawal thresholds were measured to assess perigenital sensitivity in naïve and NMS mice. A significant impact of NMS, exercise, and an NMS/exercise interaction was observed on mechanical withdrawal thresholds. Sedentary NMS males displayed significantly lower perigenital mechanical withdrawal thresholds than naïve-Sed and NMS-Ex mice. Data were log transformed prior to analysis. Bracket indicates significant effect of NMS (§, *p*=0.008), exercise (ε, *p*<0.0001), and an NMS/exercise interaction (+, *p*=0.0002), two-way ANOVA, \**p*<0.0001 vs. naïve, ^*#*^*p*<0.0001 vs. sedentary, Bonferroni posttest. n=14 (naïve-Sed, NMS-Sed), 9 (naïve-Ex), 12, (NMS-Ex).

### Voluntary wheel running normalized micturition behavior

The number and total volume of urination events were behaviorally assessed over one hour. NMS-Sed mice had a greater number of voids (Figure 3A) and larger total void volume (Figure 3B) than naïve-Sed mice. No difference in number of voids or void volume was observed between naïve-Ex and NMS-Ex mice; however, the total volume voided was significantly lower in NMS-Ex compared to NMS-Sed. These data indicate that voluntary wheel running normalized micturition behavior.

**Figure 3.**
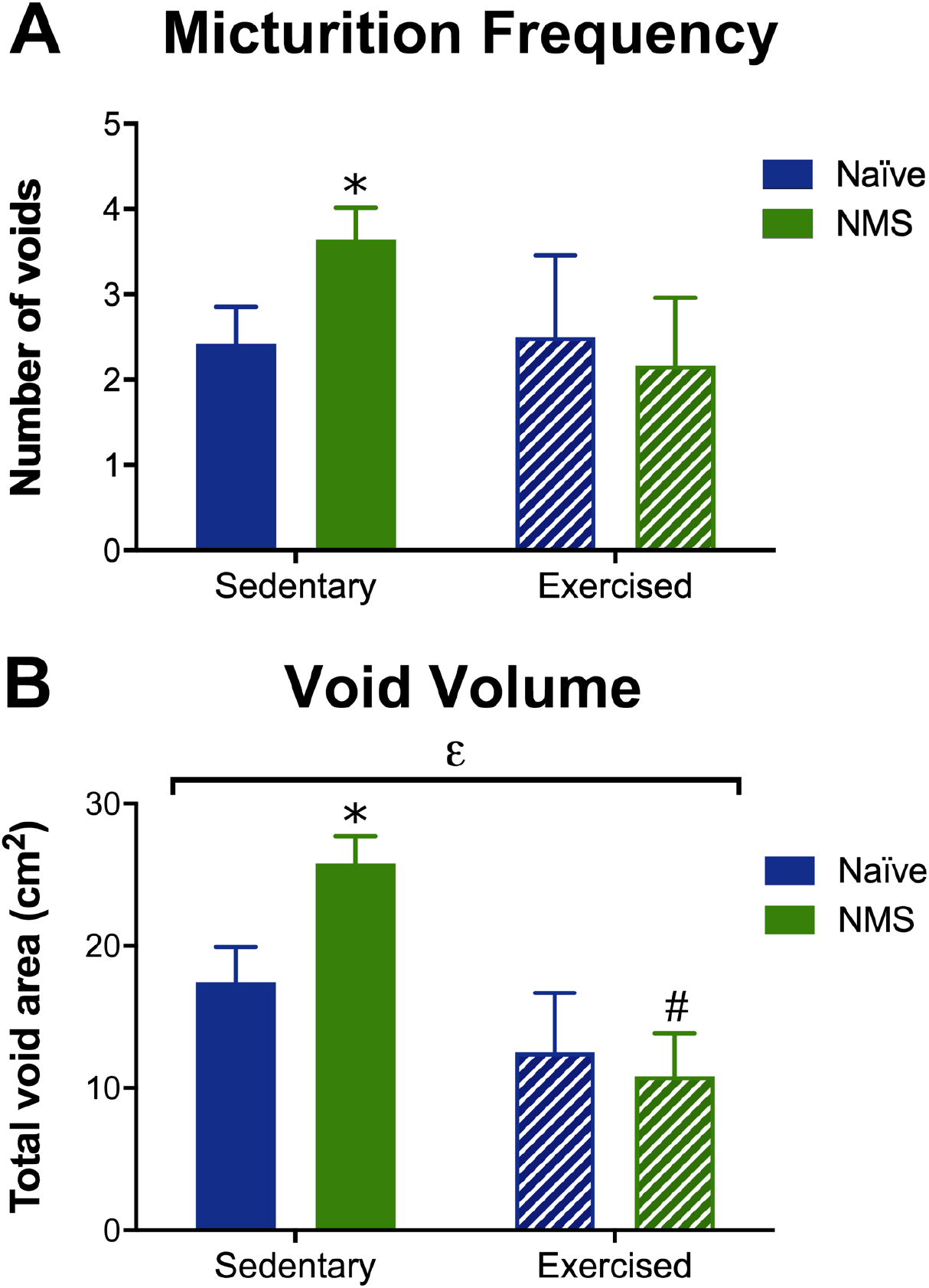
Micturition behavior was evaluated to determine changes in bladder output. **A)** The number of voids was significantly higher in NMS-Sed mice compared to naïve-Sed mice. No significant difference was observed between naïve-Ex and NMS-Ex mice. **B)** Exercise significantly decreased the total void area. The total voided volume was significantly higher in NMS-Sed mice compared to naïve-Sed and NMS-Ex. Data were rank transformed prior to statistical analysis. Bracket indicates significant effect of exercise (ε, *p*=0.0039), two-way ANOVA, \**p*<0.05 vs. naïve, ^*#*^*p*<0.01 vs. sedentary, Fisher’s LSD posttest, n=19 (naïve-Sed), 28 NMS-Sed), 6 (naïve-Ex, NMS-Ex).

### Voluntary wheel running and early life stress impact behavioral evidence of anhedonia and anxiety

Preference for 1% sucrose was measured to assess anhedonic behavior, a hallmark symptom of depression (46). We did not observe a significant effect of either NMS or exercise on the percent of 1% sucrose consumed (Figure 4A). Exercise significantly increased the volume of water consumed (Figure 4B). Total 1% sucrose consumption was significantly increased by exercise and decreased by NMS (Figure 4B), such that naïve-Ex mice consumed a significantly higher volume of sucrose than either naïve-Sed or NMS-Ex mice, indicating that NMS mice displayed anhedonic behavior regardless of voluntary wheel running status.

**Figure 4.**
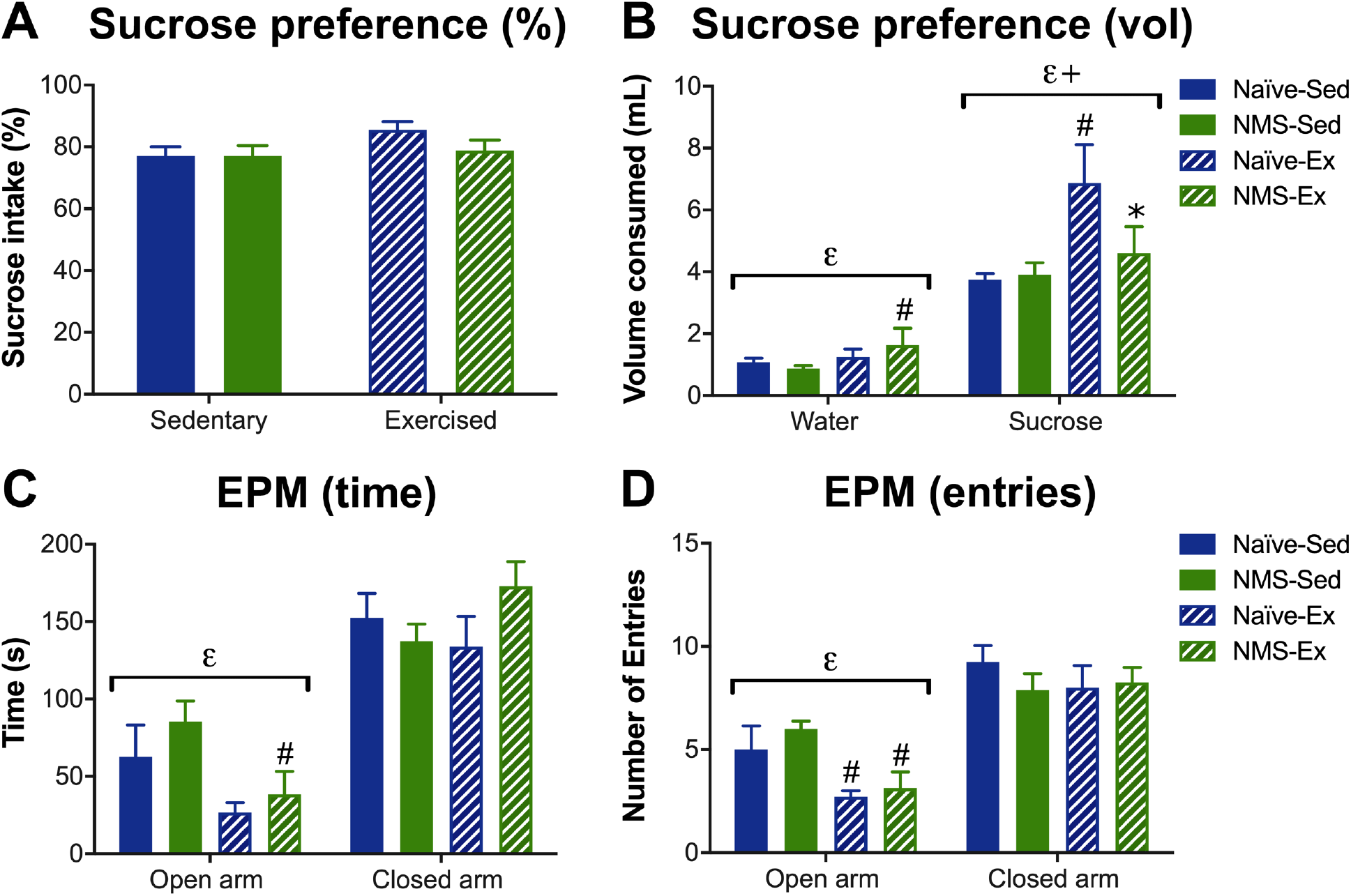
Sucrose preference and elevated plus maze testing was performed to evaluate depression- and anxiety-like behaviors, respectively. **A)** No significant impact of NMS or exercise was observed on the percent of sucrose consumed. **B)** Exercise significantly increased the volume of water (*p*=0.0346) and sucrose (*p*=0.0017) that was consumed, across naïve and NMS mice. NMS-Ex mice consumed significantly more water than NMS-Sed mice. An NMS/exercise interaction (*p*=0.0405) significantly impacted the volume of sucrose consumed, with naïve-Ex mice consuming significantly more than naïve-Sed and NMS-Ex mice. **C)** Exercise significantly decreased the amount of time spent in the open arms of an EPM, across both groups (*p*=0.0110), and particularly in NMS-Ex compared to NMS-Sed. **D)** The number of entries into the open arms was also significantly lower in exercised groups (*p*=0.0021), with naïve-Ex and NMS-Ex showing significantly fewer entries compared to their sedentary counterparts. Brackets indicate significant effect of exercise (ε) or an NMS/exercise interaction (+), two-way ANOVA, \**p*<0.05 vs. naïve, ^*#*^*p*<0.05 vs. sedentary, Fisher’s LSD posttest. **A-B**: n= 23 (naïve-Sed), 25 (NMS-Sed), 8 (naïve-Ex), 7 (NMS-Ex); **C-D**: n=8 (naïve-Sed, NMS-Sed, NMS-Ex), 7 (naïve-Ex).

An elevated plus-maze was used to assess anxiety-like behavior. Although we did not observe any significant overall effects of NMS on anxiety-like behavior, exercise significantly decreased the time spent (Figure 4C) and number of entries (Figure 4D) in the open arm of the elevated plus-maze. NMS-Ex mice spent significantly less time in the open arm (Figure 4C) and had fewer entries into the open arm (Figure 4D), compared to NMS-Sed mice, indicating increased anxiety-like behavior. No significant differences were observed between groups for measurements made regarding behavior in the closed arm.

### Voluntary wheel running reduced serum corticosterone in naïve mice and normalized mast cell degranulation rates in the bladder and prostate of NMS mice

To determine how NMS and voluntary wheel running affected downstream activation of the HPA axis, we measured serum corticosterone levels and mast cell degranulation in the bladder and prostate. Similar to our previous study (45), we observed a significant decrease in serum corticosterone in NMS-Sed mice, compared to naïve mice (Figure 5A), suggesting that NMS generates a hypocorticosteroid state in male mice. Voluntary wheel running significantly decreased serum CORT in naïve mice, but had no additional impact in NMS mice.

**Figure 5.**
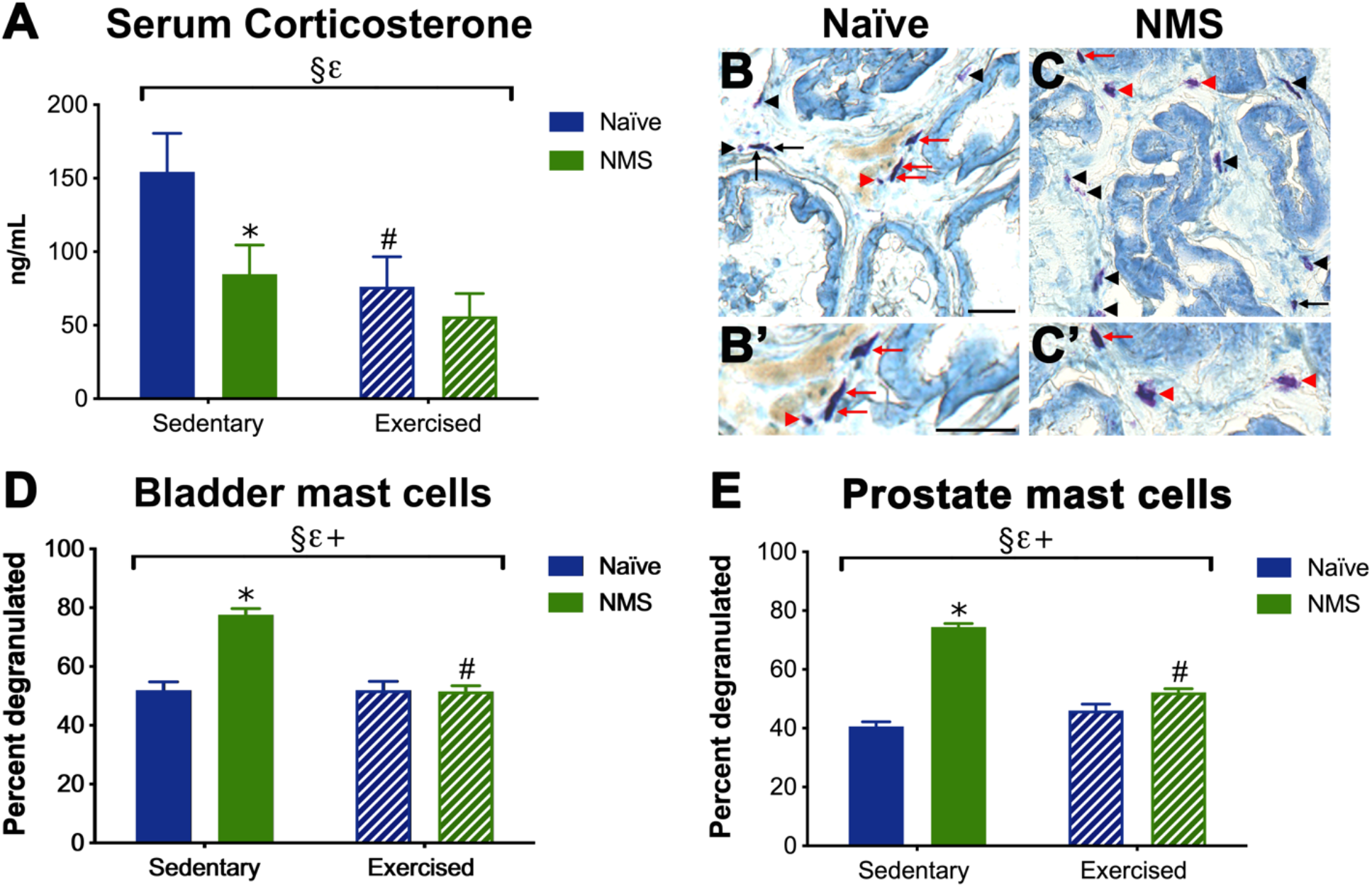
Serum corticosterone and mast cell degranulation were evaluated to determine downstream activation of the hypothalamic-pituitary-adrenal axis. **A)** NMS (*p*=0.0403) and exercise (*p*=0.0149) both significantly decreased serum CORT levels. Sedentary naïve mice had significantly higher serum corticosterone levels compared to NMS-Sed and naïve-Ex mice. Photomicrographs are shown of acidified toluidine blue-stained prostate sections from naïve **(B, B’)** and NMS **(C, C’)** mice to demonstrate intact (arrow) and degranulated (arrowhead) mast cells. Higher magnification images illustrate dense metachromasia (red arrows, **B’** and **C’**) and faint staining with extruded granules (red arrowheads, **C’**) of intact and degranulated mast cells, respectively. **D)** A significant impact of NMS (*p*<0.0001), exercise (*p*<0.0001), and an NMS/exercise interaction (*p*<0.0001) was observed on mast cell degranulation in the bladder. NMS-Sed mice exhibited significantly higher percentages of degranulated mast cells compared to both naïve-Sed and NMS-Ex mice. **E)** A significant impact of NMS (*p*<0.0001), exercise (*p*<0.0001), and an NMS/exercise interaction (*p*<0.0001) was also observed on mast cell degranulation in the prostate. NMS-Sed mice exhibited significantly higher percentages of degranulated mast cells compared to both naïve-Sed and NMS-Ex mice. Brackets indicate significant effect of NMS (§), exercise (ε), or an NMS/exercise interaction (+), two-way ANOVA; **A**) \**p*<0.05 vs. naïve, ^*#*^*p*<0.05 vs. sedentary, **C-D**) \**p*<0.0001 vs. naïve, ^*#*^*p*<0.0001 vs. sedentary, Fisher’s LSD posttest. Scale bars equal 100μm. **A**: n=10 (all groups); **D-E**: n=8 (all groups).

Acidified toluidine blue was used to visualize and quantify degranulated mast cells in cryosections of bladder and prostate tissues (Figure 5B). Consistent with our previous results (39), bladders (Figure 5D) and prostates (Figure 5B-C, E) from NMS-Sed mice contained a significantly higher percentage of degranulated mast cells, compared to tissues from naïve-Sed mice. Voluntary wheel running significantly reduced mast cell degranulation in tissues from NMS mice and had no effect in naïve mice. These data suggest voluntary running wheel activity is able to prevent increased mast cell degranulation associated with NMS.

### Voluntary wheel running increased regulatory gene expression in the hypothalamus and hippocampus

To determine how NMS and voluntary wheel running affected central brain regions involved in HPA axis activation and regulation, we measured the levels of stress-related genes in the hypothalamus, amygdala, and hippocampus using RT-PCR. CRF receptor 2 (CRF_2_) largely functions to dampen stress responses and has been shown to counter the effects of CRF_1_ signaling (47, 48). Exercise significantly increased CRF_2_ mRNA levels in the hypothalamus, and NMS-Ex hypothalamus expressed a significantly higher level of CRF_2_ mRNA compared to NMS-Sed (Table 2). The hippocampus provides negative regulation of the hypothalamus, largely through activation of glucocorticoid receptor (GR) (49). A significant interaction effect of NMS and exercise was observed on GR mRNA levels in the hippocampus (Table 2). Posthoc analyses revealed that GR mRNA levels were decreased in hippocampus from NMS-Sed mice compared to naïve-Sed mice (*p*=0.014), and were significantly increased in NMS-Ex mice, compared to NMS-Sed mice. Finally, BDNF has long been studied for its role in mediating long-term potentiation and neurogenesis in the hippocampus, particularly in response to exercise (50). Here, BDNF mRNA levels were significantly increased in the hippocampus from both NMS-Ex and naïve-Ex mice, compared to their sedentary counterparts (Table 2). Together these data suggest that exercise increases negative feedback within the hypothalamus and may increase neurogenesis in the hippocampus.

**Table 2.** Stress-related genes were evaluated in the hypothalamus, amygdala, and hippocampus using RT-PCR. Exercise had a significant impact on hypothalamic CRF_*2*_ gene expression (ε, *p*=0.0107). NMS-Ex mice had significantly higher CRF_*2*_ mRNA levels compared to NMS-Sed (^*#*^*p*=0.0482). Hippocampal GR mRNA levels had a significant NMS/exercise interaction (+, *p*=0.0216) with NMS-Ex expressing significantly higher GR mRNA than NMS-Sed (*p*=0.0143). Hippocampal BDNF mRNA levels were significantly impacted by exercise (ε, *p*=0.0461). Two-way ANOVA, Fisher’s LSD posttest. n=4 (all groups).

|  | Naïve-Sed | NMS-Sed | Naïve-Ex | NMS-Ex |
| --- | --- | --- | --- | --- |
| Hypothalamus |  |  |  |  |
| CRF | 100 ± 14.5 | 120.6 ± 3.3 | 107.7 ± 17.3 | 110.7 ± 8.7 |
| Ucn2 | 100 ± 5.7 | 106.4 ± 5.6 | 96.0 ± 22.8 | 104.8 ± 11.2 |
| CRF<sub>1</sub> | 100 ± 4.0 | 103.5 ± 10.0 | 91.5 ± 12.5 | 108.7 ± 6.3 |
| ε CRF<sub>2</sub> | 100 ± 7.5 | 162.5 ± 62.7 | 241.6 ± 66.7 | 313.1 ± 30.5<sup>#</sup> |
| GR | 100 ± 3.3 | 105.9 ± 7.5 | 97.7 ± 11.2 | 107.5 ± 5.4 |
| MR | 100 ± 7.7 | 97.0 ± 7.5 | 92.0 ± 11.7 | 97.8 ± 12.2 |
| Amygdala |  |  |  |  |
| CRF | 100 ± 14.9 | 87.0 ± 4.4 | 103.1 ± 20.0 | 102.9 ± 14.1 |
| CRF<sub>1</sub> | 100 ± 5.4 | 85.1 ± 9.9 | 97.9 ± 17.4 | 86.0 ± 13.6 |
| GR | 100 ± 1.8 | 92.5 ± 4.5 | 90.2 ± 8.5 | 88.3 ± 6.0 |
| MR | 100 ± 3.4 | 87.5 ± 6.3 | 105.2 ± 18.2 | 89.7 ± 9.6 |
| Hippocampus |  |  |  |  |
| CRF<sub>1</sub> | 100 ± 4.5 | 101.5 ± 10.5 | 105.9 ± 8.4 | 108.8 ± 3.8 |
| CRF<sub>2</sub> | 100 ± 22.1 | 108.5 ± 23.9 | 88.5 ± 16.2 | 62.0 ± 24.6 |
| + GR | 100 ± 16.6 | 65.9 ± 2.5 | 79.9 ± 15.7 | 122.5 ± 17.0<sup>#</sup> |
| MR | 100 ± 12.4 | 106.1 ± 15.7 | 99.9 ± 9.9 | 114.0 ± 12.5 |
| ε BDNF | 100 ± 8.4 | 88.0 ± 11.7 | 113.2 ± 9.2 | 113.1 ± 1.7 |

### Voluntary wheel running reverses decreased hippocampal neurogenesis in NMS mice

Following our observation of increased BDNF mRNA levels in the hippocampus of exercised mice (Table 2), we used Bromodeoxyuridine (BrdU) labeling to assess adult neurogenesis *in vivo* (Figure 6A-D) (51, 52). NMS-Sed mice had significantly fewer BrdU-labeled neurons than naïve-Sed mice (Figure 6E), suggesting a decrease in adult neurogenesis in the hippocampus. Voluntary wheel running significantly increased BrdU labeling overall, and in both naïve-Ex and NMS-Ex mice when compared to their sedentary counterparts, suggesting that exercise is capable of reversing decreased neurogenesis due to NMS.

**Figure 6.**
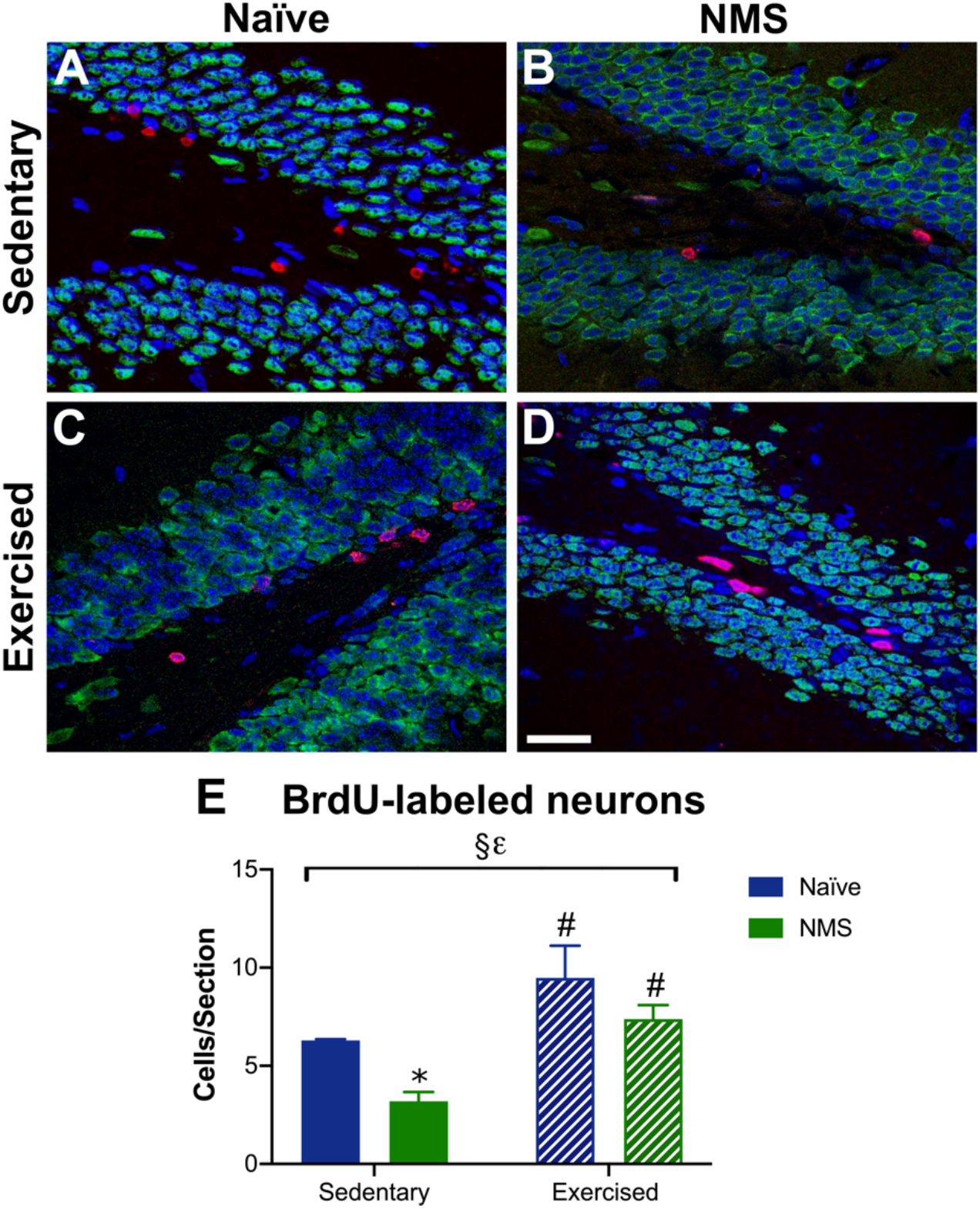
Neurogenesis was evaluated in the hippocampus using BrdU. Examples of BrdU-labeled neurons (red) in the subgranular zone of the dentate gyrus are shown from naïve-Sed (**A**), NMS-Sed (**B**), naïve-Ex (**C**), and NMS-Ex (**D**) mice. NeuN (green) labeled mature neurons in the granular zone, along with Hoechst stain (blue). **E**) A significant impact of NMS and exercise was observed on BrdU labeled neurons. NMS-Sed mice had significantly fewer BrdU-positive neurons per section, compared to naïve-Sed. Exercise significantly increased the number of BrdU-positive neurons in both naïve and NMS mice compared to their sedentary counterparts. Brackets indicate an effect of NMS (§, *p*=0.0014) or exercise (ε, *p*=0.0118); two-way ANOVA, \**p*<0.05 vs. naïve, ^*#*^*p*<0.05 vs. sedentary, Fisher’s LSD posttest. n=3 (naïve-Sed, naïve-Ex), 4 (NMS-Sed, NMS-Ex).

## Discussion

Early life stress exposure has been associated with the development of chronic pain and mood disorders later in life (3, 53–57). In patients with UCPPS, the severity of adverse childhood events is directly correlated with symptom severity and increased comorbidity, and is inversely related to likelihood of improvement (3, 10). Voluntary exercise can favorably influence neuroplasticity within the brain and improve symptoms associated with chronic pain and mood disorder (33–36, 58). Our group and others have shown that voluntary wheel running can reduce bladder hypersensitivity (38, 59) and improve obesity-related outcomes (44) in stress-induced UCPPS models, suggesting that increased physical activity may negate some of the outcomes associated with ELS. Here we provide further supporting evidence that voluntary wheel running can attenuate perigenital hypersensitivity, dysfunction, and mast cell degranulation, likely through increased hippocampal neurogenesis, which would re-establish proper negative feedback onto the HPA axis.

An important, and often overlooked, aspect of most rodent studies is the common use of sedentary caging. Rodents are not naturally sedentary and laboratory rats and mice will run up to 43 and 16km/day, respectively, if given access to a running wheel (60). Although, sedentary caging may not be a natural setting for rodents, it is likely an accurate representation of the current state of public health. Approximately 86% of the US population does not meet the current recommended guidelines for minimal physical activity (61). Men and women in non-modern cultures will average 18,000 and 14,000 steps/day, respectively, compared to 5,000 steps/day for an average modern American adult (61, 62). Sedentary behavior contributes to adult morbidity and mortality and is associated with increased depression, anxiety, and pain (63). In a meta-analysis of physical activity intervention studies in preschoolers, children, and adolescents (64), a significant association was found between increased negative affect and sedentary behavior. Although the running distances were shorter in NMS mice, compared to naïve, voluntary wheel running has consistently been sufficient to normalize most behavioral and functional outcomes in NMS mice (65), including increased adiposity on a chow diet (44).

Exercise interventions have improved pain and quality of life scores in women with chronic pelvic pain (66). Additional studies have shown similar improvements in both pain and overall well-being in irritable bowel syndrome, migraine, and fibromyalgia patients (20). Importantly, a history of being physically active, particularly having a higher level of moderate- and high-intensity physical activity, was associated with a lower risk of developing CP/CPPS in older men (67) and endometriosis in young adult women (68). Here, we report that male NMS mice display significant mechanical allodynia in the perigenital region, a finding that supports our previous research (45). Allodynia in the perigenital or pelvic region has been consistently reported in other mouse models of CP/CPPS (19, 69). Voluntary wheel running normalized mechanical withdrawal thresholds in the perigenital region of male NMS mice, consistent with the attenuation of bladder hypersensitivity in female NMS-Ex mice (38). A recent study showed that voluntary wheel running improved bladder sensory thresholds in anxiety-prone Wistar Kyoto rats that were subjected to water avoidance stress (WAS) (59), supporting the efficacy of increased physical activity for improving stress-related UCPPS-like symptoms in rodent models.

Increased urinary urgency and frequency are characteristic symptoms of IC/PBS and, in some cases, CP/CPPS. Mouse models of prostatic inflammation have demonstrated increased micturition frequency (69, 70), supporting the symptomatic overlap between prostate and bladder disease. We report here that voluntary wheel running prevented the increase in both void frequency and total urine output in sedentary male NMS mice. Preclinical models of IC/PBS and diabetes have shown improvement in bladder function following exercise. The previously mentioned study on voluntary wheel running in WAS-exposed Wistar Kyoto rats also demonstrated a significant decrease in void frequency, compared to non-exercised rats, as well as increased bladder capacity and threshold pressure (59). Interestingly, the authors reported a significant correlation between distance ran and void frequency, such that the rats that ran the greatest distance also displayed the largest decrease in voids, suggesting a dose response effect of exercise on bladder function. A study using forced wheel running in the db/db mouse model of type 1 diabetes showed improvement in voiding frequency and urodynamic parameters (71), suggesting that improvements in bladder function due to exercise are not dependent upon the underlying disease or disorder.

Increasing physical activity through exercise interventions has been shown to reduce perceptions of stress and decrease depression and anxiety (72). Voluntary wheel running is naturally rewarding to rodents and can be used as a read-out of general well-being (73), suggesting that the significantly lower running distance of NMS mice may be indicative of impaired mood. Sucrose preference is commonly used to measure reward-seeking behavior and has been validated as a biobehavioral measure of anhedonia (46). Although we did not observe a significant effect of NMS on sucrose preference, the amount of sucrose consumed was increased in naïve-Ex mice and not in NMS-Ex mice, suggesting that exercise had a diminished impact on NMS mice. A recent systematic review and meta-analysis on anxiety-like behaviors in rodent models of NMS revealed increased defensive behavior in rats, but not in mice (74). One of the analyzed tasks included elevated plus maze, which measures exploratory-defense behaviors (42). In concordance with this meta-analysis, we did not observe any effects of NMS on EPM behaviors. Surprisingly, voluntary wheel running decreased both the amount of time spent and the number of entries into the open arms of an elevated plus maze, indicating an increase in anxiety-like behavior.

Altered regulation and output of the HPA axis has been implicated in multiple chronic pain disorders, including those comprising UCPPS (20). Both hypercortisolism and hypocortisolism have been reported in adults with a history of childhood abuse or stress, the latter being postulated to arise as a compensatory response to a period of hypercortisolism and excess glucocorticoid release (75). Here, we report that male NMS-Sed mice have a significantly lower serum CORT level than naïve-Sed mice. We have previously reported lower CORT levels in male NMS mice (45), but increased CORT levels in female NMS mice, which were decreased following a similar voluntary wheel running intervention (38). In this study, voluntary wheel running significantly reduced serum CORT in naïve mice and had relatively little impact on CORT levels in NMS mice. Despite having relatively low serum CORT levels, NMS-Sed mice had significantly higher levels of degranulated mast cells in both the bladder and prostate. Similar histological findings, and molecular evidence of mast cell activation, have been reported for patients with UCPPS (18–20). Voluntary wheel running completely attenuated the increase in degranulated mast cells, suggesting a normalization of HPA axis output. Further investigation into peripheral CORT and CRF signaling is needed to fully understand the mechanism underlying the response of mast cells to voluntary exercise.

The hippocampus is particularly sensitive to excess glucocorticoids during early development and overexposure can have permanent effects on hippocampal structure, function, and gene expression (76). Human studies have shown that exposure to ELS is associated with decreased hippocampal gray matter volume (11–16) and epigenetic modifications on stress-responsive genes (77, 78). Voluntary wheel running has a significant impact on increasing hippocampal neurogenesis and BDNF expression, and preventing depression-like behaviors in rodents (33–36). These outcomes are often not observed following forced exercise interventions, such as treadmill running, due to stress-induced effects (35, 36). Here, we saw a significant impact of voluntary wheel running on BDNF expression in the hippocampus and an interaction effect of NMS and exercise on GR expression, such that it significantly increased GR mRNA levels in NMS-Ex mice compared to NMS-Sed mice. We also observed a significant increase in CRF_2_ mRNA levels in hypothalamus from NMS-Ex mice, compared to NMS-Sed mice, all of which likely reflects increased negative regulation within and upon the HPA axis. In addition to increased BDNF mRNA levels, we also observed a significant increase in BrdU-labeled neurons in the dentate gyrus of the hippocampus of both naïve- and NMS-Ex mice, compared to their sedentary counterparts. The NMS-Sed mice had a significant decrease in BrdU-labeled neurons, potentially reflecting the clinical observations of decreased gray matter volume in patients that had been exposed to ELS (15, 16).

In conclusion, our data show that early life stress exposure, here in the form of NMS, increases urogenital sensitivity, dysfunction, and mast cell degranulation, likely due to altered stress-related gene expression within and diminished negative feedback onto the HPA axis. Voluntary wheel running, although lower in NMS mice, was sufficient to normalize perigenital sensitivity, micturition behavior, and mast cell degranulation rates. Exercise significantly increased regulatory gene expression in the hippocampus, as well as increased neurogenesis in the dentate gyrus. These findings support evidence in both preclinical and clinical literature (20, 24, 28, 29, 67) suggesting that exercise may provide a powerful non-pharmacological therapeutic option for the treatment of chronic pain disorders, particularly in patients with comorbid mood disorders and/or a history of early life stress.

## Author contributions

IMF, DEW, and JAC designed the research study; IMF, BMJ, ADB, RMS, ANP, and OCE performed the experiments; IMF, BMJ, ANP, OCE, and JAC analyzed the data; IMF and JAC wrote the manuscript.

## Acknowledgments

We would like to thank Ruipeng Wang and Janelle Ryals for technical assistance, and Drs. Paige Geiger, Tomas Griebling, and Kenneth McCarson for expert advice and feedback on experimental design, methodology, and interpretation. We acknowledge that part of this work appears in dissertation form.

